# A thymus-independent artificial organoid system supports complete thymopoiesis from Rhesus macaque-derived hematopoietic stem and progenitor cells

**DOI:** 10.1101/2025.08.29.673172

**Authors:** Callie Wilde, Saleem Anwar, Yu-Tim Yau, Sunil Badve, Yesim Gokmen Polar, John D. Roback, Rama Rao Amara, R. Paul Johnson, Sheikh Abdul Rahman

## Abstract

Thymic output has been extensively studied. While advanced ex-vivo T cell generative approaches exist for mouse and human models, such advancements for nonhuman primate model are lacking. We report the establishment of a rhesus macaque-specific artificial thymic organoid (Rh-ATO) system enabling robust ex vivo T cell generation from CD34⁺ hematopoietic stem and progenitor cells (HSPCs). A continuum of distinct thymopoietic stages were recorded - robust T cell specification of HSPCs resembling thymus seeding progenitors, emergence of CD4+CD3-immature single positive, CD4+CD8+ double positive thymocytes, and finally, generation of CD4⁺ and CD8⁺ single-positive T cell subsets expressing CD38, consistent with recent thymic emigrant phenotype. These events closely mirrored Bonafide thymopoietic stages observed in the thymus. T cells expressed TCRs and exhibited polyfunctional cytokine expression. Thus, we report first demonstration of an off the shelf NHP-specific 3D system recapitulating thymopoiesis, providing a translational platform for modeling T cell development, therapeutic strategies, and immunopathogenesis.

## Introduction

T cell development involves a stepwise transformation of hematopoietic stem and progenitor cells (HSPCs), which undergo initial T lineage commitment in the bone marrow to generate thymus-seeding progenitors (TSPs) (Cordes et al., 2022; Hoebeke et al., 2007). Upon thymic entry, TSPs-typically CD3^-^CD4⁻CD8⁻CD34⁺CD38^⁻/+^CD7^⁻/+^-upregulate markers such as CD7 and the T cell factor 1 (TCF-1), differentiating into immature single-positive (ISP) thymocytes (Ashby and Hogquist, 2023; Germain, 2002; Hoebeke *et al*., 2007; La Motte-Mohs et al., 2005; Six et al., 2007). ISPs express CD4 but lack CD3, and transition into CD4⁺CD8⁺ double-positive (DP) thymocytes, which subsequently upregulate CD3. Positive selection via MHC-I or MHC-II leads to the emergence of CD8⁺ and CD4⁺ single-positive (SP) T cells, respectively. The SP CD8 and CD4 T cells leave thymus into the circulation, where they are called recent thymic emigrants and express high levels of surface CD38 (Bohacova et al., 2024). Accurately modeling this ontogeny ex vivo is critical not only for generating therapeutic T cells but also for advancing our understanding of T cell regenerative biology and thymic disruption in disease.

Nonhuman primates (NHPs) offer a physiologically relevant preclinical model for studying T cell development in the context of chronic and infectious diseases (Estes et al., 2018; Tarantal et al., 2022). Prior attempts to model thymopoiesis from NHP-derived HSPCs demonstrated T cell selection and development (Rosenzweig et al., 1996a; Rosenzweig et al., 1996b; Rosenzweig et al., 2001). These systems required co-culturing HSPCs with either rhesus or porcine-derived primary thymus stromal cells. Similar strategies in human systems employed CD34⁺ HSPCs with thymic epithelial components (Akkina et al., 1994; Jenkinson et al., 1982; Peault et al., 1991; Rosenzweig *et al*., 1996b; Rosenzweig *et al*., 2001; Yeoman et al., 1993), however broader adoption has been limited by the need for primary thymic tissue and its short shelf life. To overcome these limitations, murine stromal cell lines (OP9 or MS-5) were engineered to express Notch ligands DLL1 or DLL4, enabling thymus-independent co-culture systems (Kutlesa et al., 2009; La Motte-Mohs *et al*., 2005; Mohtashami et al., 2023; Mohtashami et al., 2013; Schmitt and Zuniga-Pflucker, 2002). When co-cultured with HSPCs from human, NHP, or mouse origins, these systems biased differentiation toward the T cell lineage, in contrast to the default B or NK pathways (Awong et al., 2008; De Smedt et al., 2004; Mohtashami *et al*., 2023; Radtke et al., 2017) underscoring the significance of notch signaling.

Despite conserved notch pathway, human or NHP HSPCs cultured with OP9/MS5-DLL1/4 system have shown inefficient selection of T cell precursors leading to fewer productively TCR rearranged T cell subset development (Awong et al., 2011; Chung et al., 2014; De Smedt *et al*., 2004; La Motte-Mohs *et al*., 2005; Mohtashami *et al*., 2023; Radtke *et al*., 2017; Van Coppernolle et al., 2009). Improved outcomes were observed when human or mouse HSPCs were aggregated with stromal cells expressing species-specific DLL1 in 3D culture (Montel-Hagen et al., 2022; Seet et al., 2017). These so called artificial thymic organoid systems generated consistent T cell developmental outcomes with enhanced TCR^+^ T cell selection. Nonetheless, studies testing ex-vivo T cell generation methods in NHP model is limiting (Radtke *et al*., 2017; Rosenzweig *et al*., 1996b). The field lacks a validated, species-specific 3D organoid system for NHPs capable of supporting functional T cell output.

Here, we report the development and validation of a rhesus macaque-specific artificial thymic organoid (Rh-ATO) system that supports efficient ex vivo thymopoiesis from CD34⁺CD3⁻ bone marrow-derived HSPCs. Using MS5 stromal cells engineered to express RM-specific DLL1 and aggregated in 3D cultures with RM-derived progenitors, the Rh-ATO platform recapitulated sequential thymic intermediates—including CD7⁺ TSPs, CD4⁺CD3⁻ ISPs, CD4⁺CD8⁺ DPs-and yielded mature CD3⁺TCR⁺ SP CD4⁺ and CD8⁺ T cell subsets expressing CD38, resembling recent thymic emigrants. Notably, the system also supported robust T cell differentiation from human HSPCs, underscoring its translational potential. This study establishes a Rh-ATO as the first NHP-specific, thymus-independent organoid system capable of generating functional T cells ex vivo.

## Results

### Generation and characterization of MS5 stromal cells expressing Rhesus macaque (*Macaca mulatta*) DLL1

The murine bone marrow stromal cell line MS5 was genetically engineered to constitutively express rhesus macaque Delta-like canonical Notch ligand 1 (Rh-DLL1). To achieve stable expression, a lentiviral construct was designed encoding DLL1, an enhanced green fluorescent protein (eGFP) reporter, and a puromycin resistance selection cassette was designed for genomic integration (Fig. 1A). Flow cytometric analysis confirmed efficient transduction, with more than 99% of MS5 cells co-expressing eGFP and Rh-DLL1 (Fig. 1B-C). To eliminate non-transduced cells and enrich for DLL1-expressing populations, cultures were subjected to puromycin selection, resulting in exclusive survival of cells harboring the Rh-DLL1 cassette (Fig. 1D). The MS5-Rh-DLL1 was free of any potential mycoplasma contamination (Fig. 1E). The resulting MS5-Rh-DLL1 stromal cells were used to establish an Rh-ATO as described in the method section.

**Figure 1.**
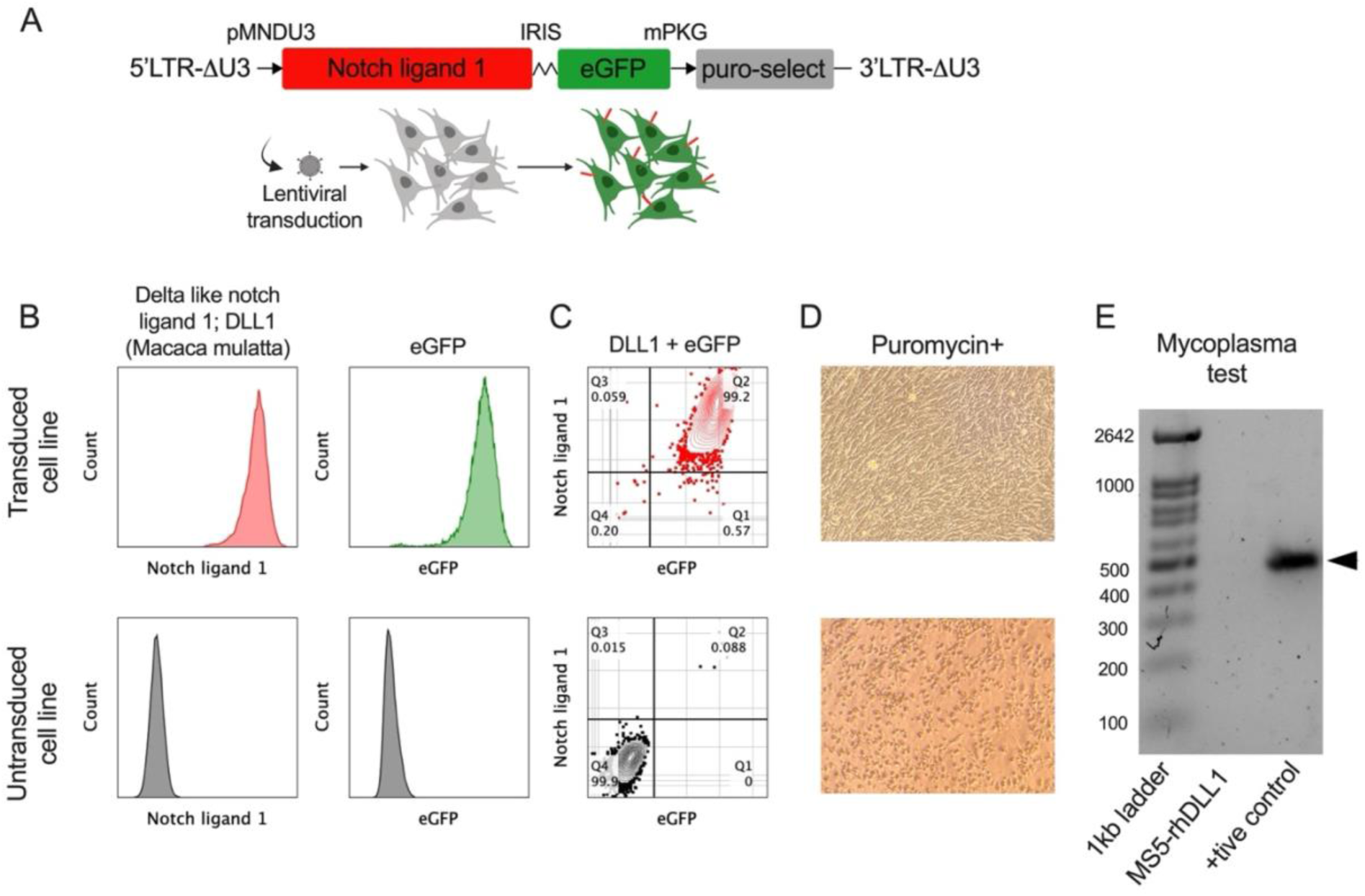
Characterization of a recombinant MS-5 stromal cell line expressing *Macaca mulatta* notch ligand 1 gene (A-D). (**A**) Cartoon showing lentiviral construct design with *Macaca mulatta* Notch ligand 1, eGFP reporter, and puromycin coding sequences. (**B, C**) Flow cytometry plots showing positive surface expression of Notch ligand 1, eGFP (**B,** upper plots in red and green respectively), and their negative expressions in the non-transduced cell line (**B**, lower plots in grey). Flow cytometry plots showing positive (**C,** upper plot in red) and negative (**C,** lower plot in grey) co-expression of DL1 and eGFP in the transduced and non-transduced cell line respectively. (**D**) Widefield image taken by iPhone mobile attachment at 5X showing viable transduced cell line (**D**, upper image) and dead non-transduced cells (**D**, lower image) cultured in the presence of 5ug/mL puromycin. (E) Gel electrophoresis showing absence of mycoplasma in the newly established MS5-Rh-DLL1 cell line.

### The Rh-ATO system supported T cell–biased differentiation of rhesus CD34⁺ HSPCs

To evaluate which lymphoid lineages the Rh-ATO system generated following differentiation of CD34+ HSPCs, we aggregated CD3⁻CD34⁺ HSPCs isolated from rhesus macaque bone marrow with MS5-Rh-DLL1 stromal cells forming 3D structures (Fig. 2A). Common lymphoid lineage (T, NK, and B-cell) differentiation of HSPCs was analyzed at week four of culture by flow cytometry (Fig. 2B). Rh-ATO cultures demonstrated a pronounced bias toward T cell (CD3⁺) differentiation, as reflected by significantly higher CD3⁺ T cell lineage (geomean frequency ∼24%) as a frequency of total CD45⁺ lymphocytes compared with B cells (geomean frequency ∼0.8%) and NK cells (geomean frequency 0.2%) (Fig. 2C). Further phenotypic profiling revealed marked enrichment of CD7⁺ cells (geomean frequency ∼80%)—a hallmark of thymocyte differentiation. CD7+ cells showed predominant bias toward T cell lineage and not B- or NK cells differentiation (Fig. 2D), further confirming T cell lineage bias. These findings demonstrated that the Rh-ATO recapitulates thymic cues necessary for T lineage specification from RM CD34⁺ HSPCs over other lymphoid lineages.

**Figure 2.**
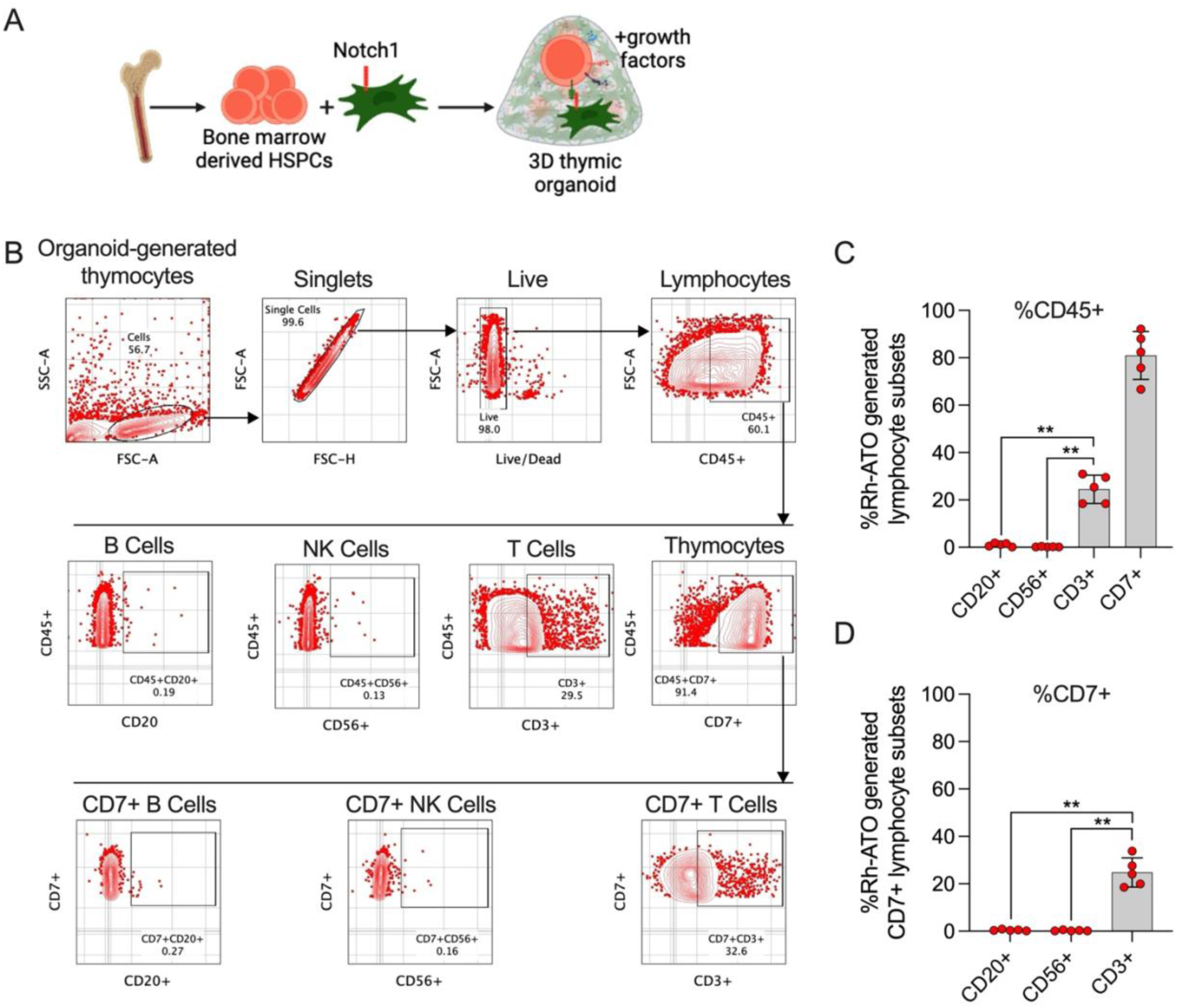
Establishment of a thymus-independent artificial organoid (Rh-ATO) system using nonhuman primate bone marrow-derived hematopoietic stem and progenitor cells (A-D). (**A**) Cartoon showing Rh-ATO establishment scheme. (**B**) Gating strategy showing flow cytometry analysis of distinct lymphocyte lineages (B, NK, and T cells) generated by Rh-ATO system at week 4 of culture. (**C**) Bar graph with individual data points showing frequencies of B cells (CD20+), NK cells (CD56+), T cells (CD3+), and thymocytes (CD7+) as a frequency of total lymphocytes (CD45+). (**D**) Bar graph with individual data points showing frequency of B cells (CD20+), NK Cells (CD56+), and T cells (CD3+), as frequency of thymocyte (CD7+) population. <u>Experimental replicates:</u> A total of five organoids (N = 5) were analyzed; 2 technical organoid replicates from two independent experiments and 1 organoid from a third independent experiment were analyzed. Statistical significance was calculated by Mann-Whitney U test. *P < 0.05.

### Rh-ATO recapitulated early T cell lineage specification and generated terminally differentiated T cells resembling recent thymic emigrants

We next examined whether the rhesus artificial thymic organoid (Rh-ATO) system supports early T cell lineage specification and the emergence of terminally differentiated T cells resembling recent thymic emigrants (RTEs). To define early progenitor phenotypes, we first analyzed CD34⁺CD4⁻CD8⁻ double-negative hematopoietic progenitors in the thymus and peripheral blood (Fig. 3A-B). As expected, the thymus contained a markedly higher frequency of CD34⁺CD7⁺ thymus-seeding progenitors (TSPs) (geomean ∼66%) compared to blood (geomean ∼1%). Consistent with this pattern, Rh-ATO cultures markedly induced CD7 expression on CD34⁺ progenitors, achieving frequencies comparable to thymic TSPs. Over 28 days of culture, CD34⁺CD7⁺ cells expanded from ∼18% at day 0 to ∼62% at day 28, indicating efficient T cell lineage priming within the organoid microenvironment. The frequency of CD7⁺CD34⁺ cells in Rh-ATO cultures remained significantly higher than that observed in blood, mirroring thymic conditions.

**Figure 3.**
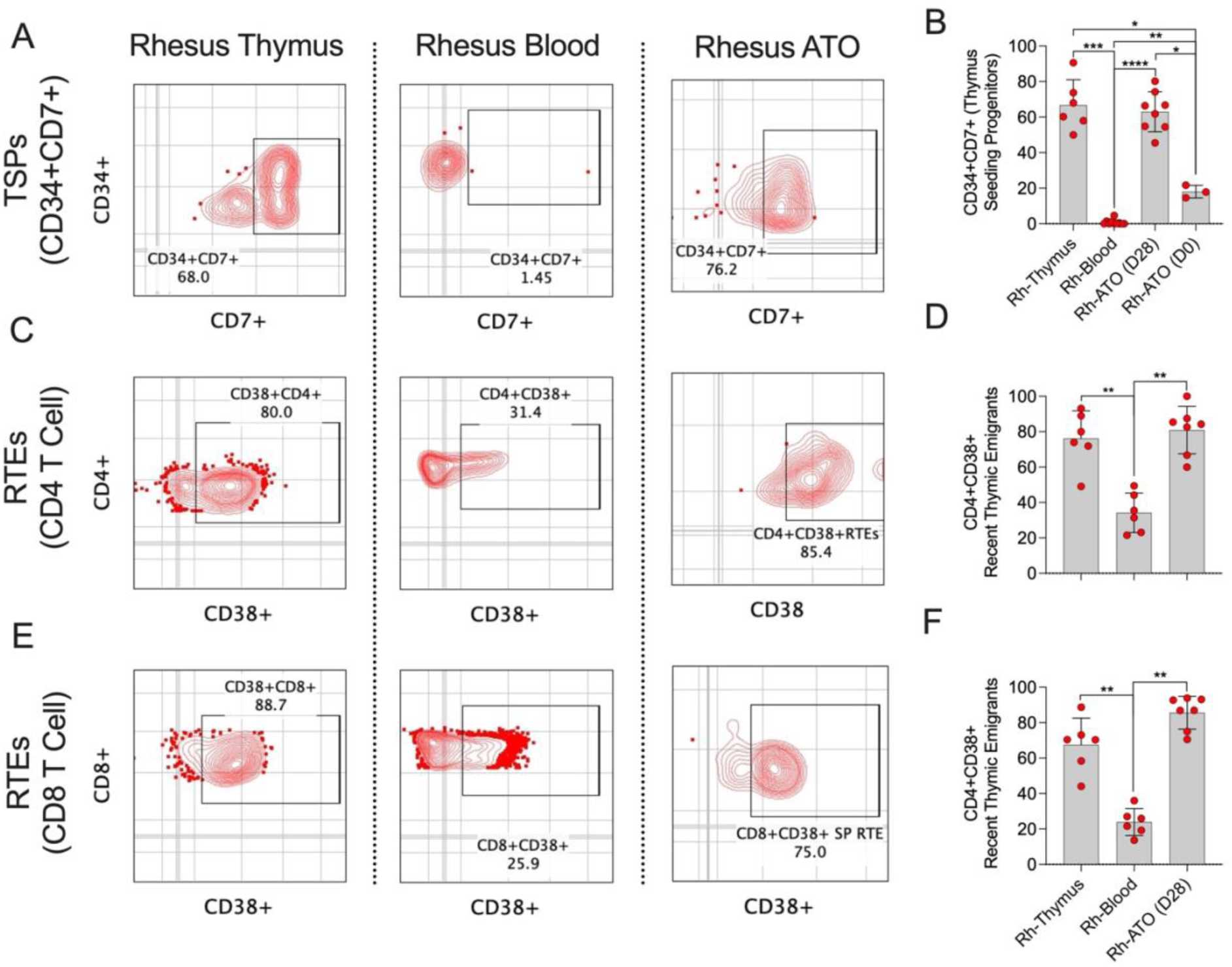
Rh-ATO generated thymocytes exhibited features of both thymus seeding progenitors (TSPs) and recent thymic emigrants (RTEs) (A-F). (**A, B**) Flow cytometry plots showing frequencies of TSPs expressing CD7 on CD4-CD8-double negative CD34+ HSPCs in rhesus macaque derived thymus, blood, and Rh-ATO. (**B**) Bar graph with individual data points is shown. (**C-F**) Flow cytometry plots and bar graphs with individual data points showing frequencies of RTEs marked by the expression of CD38 on CD4+ (**C, D**) and CD8+ (**E, F**) T cell subsets in rhesus macaque derived thymus, blood, and Rh-ATO. <u>Experimental replicates:</u> Thymus and blood samples were obtained from six Rhesus macaques (N = 6). For data in Figure 3B, 2 technical organoid replicates from four independent experiments (N = 8 organoids) were analyzed at day 28 of culture, and three organoids (N = 3) from three independent experiments were analyzed at day 0 of culture. For data in Figure 3D-F, seven organoids (N = 7) from above experiments were additionally analyzed for RTEs. Statistical significance was calculated by Mann-Whitney U test.

We next assessed whether Rh-ATO–derived mature T cells exhibited phenotypic features of RTEs, defined by CD38 expression on CD4⁺ and CD8⁺ T cell subsets (Fig. 3C-F). CD38⁺CD4⁺ T cells generated in Rh-ATOs reached ∼80%, closely matching thymic RTE frequencies (∼75%) and substantially exceeding those in blood (∼32%). Similarly, CD38⁺CD8⁺ T cells in Rh-ATOs (∼85%) paralleled thymic frequencies (∼85%) and were significantly enriched compared to blood (∼24%). These data demonstrate that the Rh-ATO system not only drives early T cell specification from hematopoietic progenitors but also supports terminal differentiation into T cells phenotypically similar to those recently emigrated from the thymus.

### Rh-ATO system recapitulated canonical thymic intermediates within 28 days

To verify that the Rh-ATO system recapitulated authentic thymic development, we first analyzed thymocytes directly isolated from rhesus macaque (RM) thymus. As anticipated, phenotypic analysis revealed the canonical thymocyte stages, including CD4⁺CD3⁻ immature single-positive (ISP) cells, CD4⁺CD8⁺ double-positive (DP) intermediates, and terminally differentiated CD3⁺ single-positive T cells (Fig. 4A). We next examined whether Rh-ATO–derived T cell development follows this trajectory by longitudinally tracking cultures over four weeks (Fig. 4B-G). At day 7, ISPs dominated the population (range 30–58%; geomean 39%), with minimal DP representation (∼3%). By day 14, DP cells expanded markedly (geomean 10%), accompanied by decreased ISP frequencies (∼23%) and the first appearance of CD3 expression (∼2%). At this stage, CD4⁺ (∼8%) and CD8⁺ (∼6%) single-positive cells also emerged. By day 28, cultures contained robust populations of DP cells (geomean 36%) and mature CD3⁺ T cells (geomean 29%), including CD4⁺ (∼7%) and CD8⁺ (∼34%) single-positive subsets, while ISP frequencies further declined (∼21%). This ISP>DP>SP progression closely mirrored in vivo thymopoiesis.

**Figure 4.**
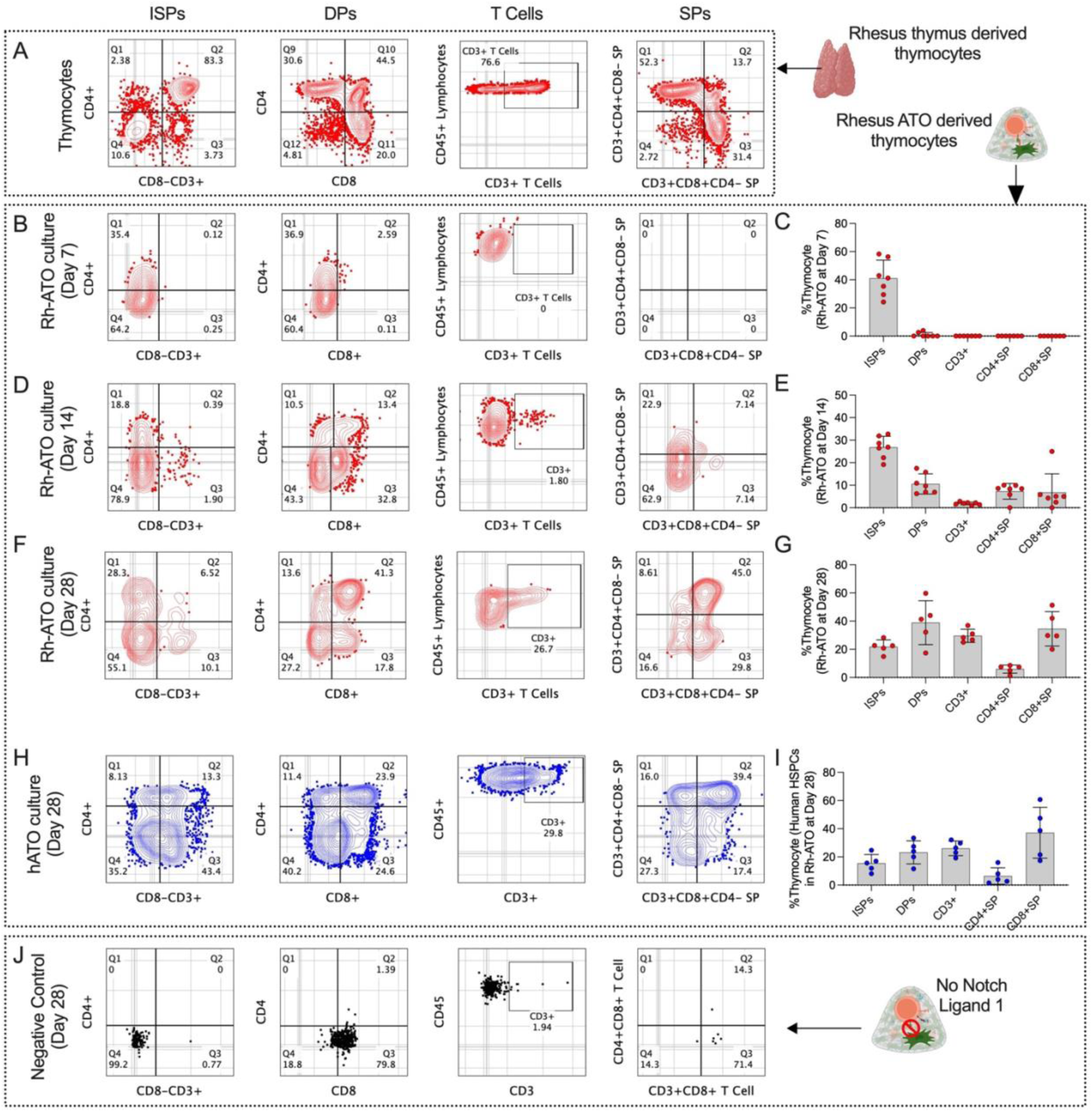
Characterization of distinct thymic intermediate stages generated under Rh-ATO system (A-J). (**A**) Flow cytometry plots showing established Bonafide thymocyte intermediate populations in rhesus macaque thymus - CD4+CD3-immature single positive (ISPs), CD4+CD8+ double positive (DP), CD3+ T cell, CD3+CD4+CD8-, and CD3+CD8+CD4-single positive (SP) T cells. (**B-G**) Flow cytometry plots and bar graphs with individual data points showing developmental kinetics of distinct thymic intermediate population stages at day 7 (**B, C**), day 14 (**D, E**), and day 28 (**F, G**) of Rh-ATO culture establishment using rhesus macaque bone marrow-derived HSPCs. (**H, I**) Flow cytometry plots and bar graphs with individual data points showing generation of distinct thymic populations at day 28 of Rh-ATO culture establishment using mobilized human peripheral blood derived HSPCs. (**J**) Flow cytometry plots showing inability of thymocyte generation of HSPCs in Rh-ATO lacking Rh-DLL1 expression. <u>Experimental</u> <u>replicates:</u> For data in Figure 4C-E, a total of seven organoids (N = 7) were analyzed from three independent experiments. For data in Figure 4G-I, a total of five organoids (N = 5) were analyzed from three independent experiments. Statistical significance was calculated by Mann-Whitney U test. *P < 0.05.

To evaluate cross-species applicability, we aggregated human mobilized peripheral blood CD34⁺ HSPCs with Rh-DLL1–expressing stromal cells and analyzed thymopoiesis at week 4 (Fig. 4H-I). Human HSPCs generated comparable intermediates and mature T cell subsets, including ISPs (15%), DP cells (22%), and CD3⁺ T cells (26%), with CD4⁺ (∼5%) and CD8⁺ (∼33%) single-positive populations resembling RM-derived outcomes. In contrast, organoids lacking Rh-DLL1 failed to produce T cells, underscoring the essential role of Notch signaling in driving T lineage specification (Fig. 4J). Together, these findings demonstrate that the Rh-ATO platform robustly supports canonical thymic development from both RM and human HSPCs, enabling faithful modeling of T cell ontogeny ex vivo.

### Rh-ATO derived T cells expressed T-cell receptor and demonstrated polyfunctional cytokine response upon stimulation

To evaluate whether T cells generated in the Rh-ATO system undergo productive TCR rearrangement, we first assessed expression of TCRαβ, the predominant TCR subtype. At four weeks of culture, ∼15% of CD3⁺ T cells expressed TCRαβ (Fig. 5A), with both CD4⁺ and CD8⁺ single-positive subsets showing comparable TCRαβ expression (Fig. 5B-C). Next, we examined the functional capacity of Rh-ATO–derived T cells by quantifying cytokine responses following stimulation. Enriched CD3⁺ T cells stimulated with PMA/ionomycin (PMAi) exhibited robust induction of IFN-γ, TNF-α, and IL-2 compared to unstimulated controls (Fig. 5D-I). Boolean analysis revealed that 54% of T cells produced at least one cytokine (single cytokine responders; ∼4–15% for individual cytokines), 37% co-expressed two cytokines, and 9% co-expressed all three cytokines, indicating a polyfunctional response profile (Fig. 5F-I). Collectively, these results demonstrated that Rh-ATO-generated T cells acquire productive TCRαβ expression and develop polyfunctional effector capacity, closely mirroring thymus-derived T cell function and confirming the fidelity of the Rh-ATO platform for modeling T cell development.

**Figure 5.**
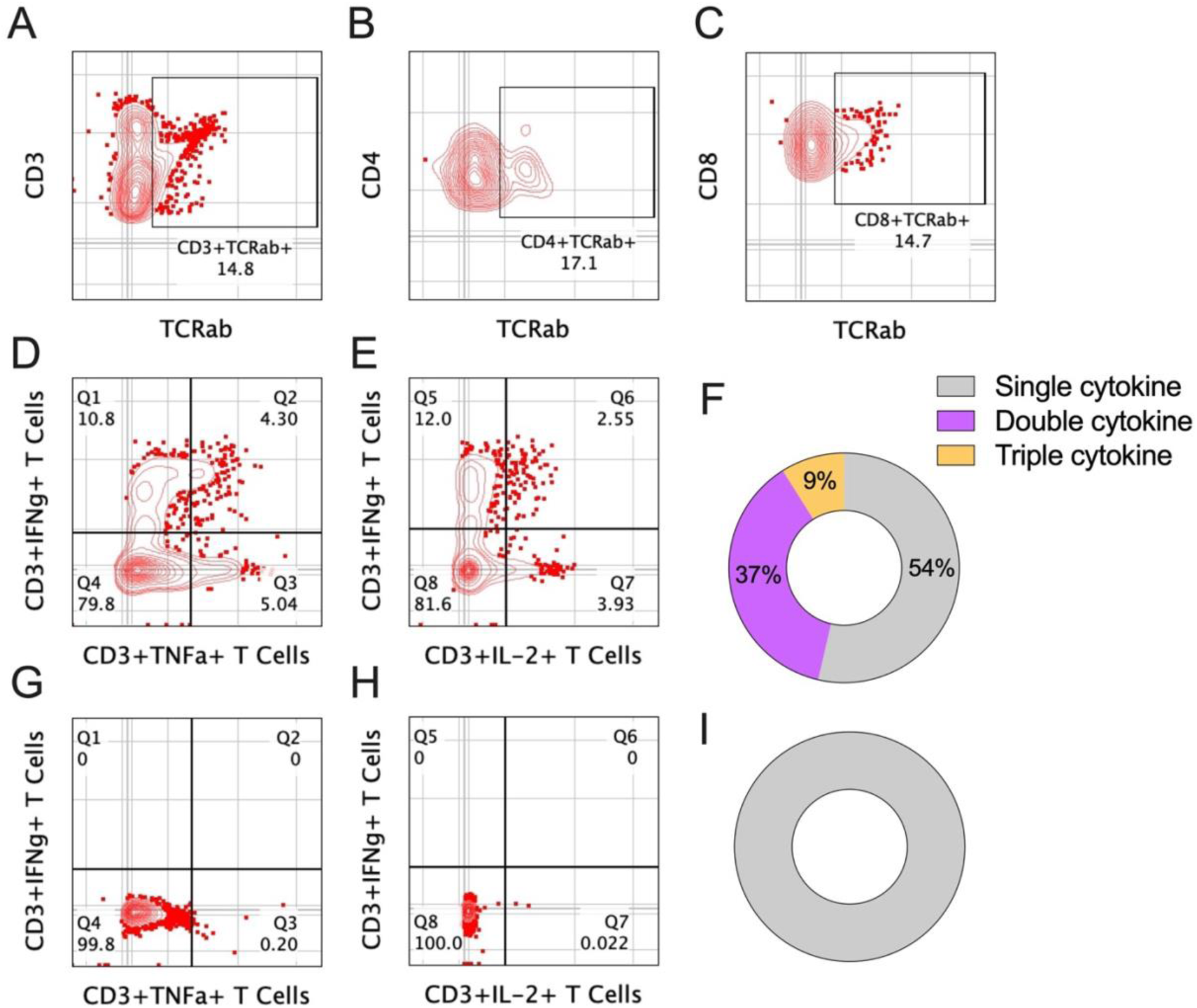
T cells generated in Rh-ATO expressed T-cell receptor and co-expressed cytokines upon stimulation (A-F). (**A-C**) Flow cytometry plots showing expression of TCRαβ of CD3+ (**A**), CD4+ (**B**), and CD8+ (**C**) T cell subsets. (**D-I**) Flow cytometry plots showing indicated cytokine expression by T cells either upon stimulation with phorbol 12-myristate-13-acetate (PMA), and ionomycin (i) (**D, E**), or no stimulation (**G, H**). Rh-ATO (N = 6) generated, and enriched T cells expressing IFNγ^+^, TNFα^+^, and IL2^+^ cytokines. Boolean analysis of cytokine co-expression is shown for PMAi stimulated (**D, E**) and unstimulated (**G, H**) T cells. Single cytokine = IFNγ^+^, TNFα^+^, and IL2^+^; Double cytokine = co-expression of any following combinations: IFNγ^+^/TNFα^+^, IFNγ^+^/IL2^+^, TNFα^+^/IL2^+^; and Triple cytokine = co-expression of IFNγ^+^/TNFα^+^/IL2^+^ cytokines. A total of 6 Rh-ATO were pooled to enrich CD3+ T cells for cytokine analysis.

## Discussion

In this study, we describe the development and characterization of a rhesus macaque–specific artificial thymic organoid (Rh-ATO) system that supports efficient ex vivo T cell differentiation from bone marrow–derived hematopoietic stem and progenitor cells (HSPCs). The Rh-ATO recapitulated key stages of thymopoiesis, including the emergence of CD7⁺CD34⁺ double-negative progenitors resembling thymus-seeding progenitors (TSPs), progression to immature single-positive (ISP) and double-positive (DP) thymocytes, and maturation into CD4⁺ and CD8⁺ single-positive T cells expressing functional TCRs. Importantly, these T cells exhibited polyfunctional cytokine responses, indicating acquisition of immunological competence. To our knowledge, this represents the first nonhuman primate (NHP)–specific 3D organoid platform that faithfully reproduced canonical thymic development.

Previous systems for ex vivo T cell generation relied largely on murine stromal cell lines expressing DLL1 or DLL4 (e.g., OP9-DLL1, MS5-DLL1) and have been instrumental in directing both murine and human HSPCs toward the T cell lineage (Chung *et al*., 2014; de Pooter and Zuniga-Pflucker, 2007; De Smedt *et al*., 2004; Kutlesa *et al*., 2009; La Motte-Mohs *et al*., 2005; Mohtashami *et al*., 2023; Mohtashami *et al*., 2013; Schmitt and Zuniga-Pflucker, 2002; Van Coppernolle *et al*., 2009). However, these approaches frequently yielded suboptimal CD8⁺ T cell output and required prolonged culture periods. Advances in 3D organoid culture have underscored the importance of spatial organization for efficient thymocyte selection and maturation (Montel-Hagen *et al*., 2022; Poznansky et al., 2000; Seet *et al*., 2017). Our findings extend this concept to the NHP setting by demonstrating that a rhesus-specific DLL1-expressing MS5 cell line can drive robust T cell development in a 3D configuration. Compared with human DLL1-based ATOs, the Rh-ATO achieved higher frequencies of mature CD4⁺ (∼5%) and CD8⁺ (∼32%) T cells within four weeks of culture and exhibited more rapid organoid compaction (2–5 days vs. 1–2 weeks). The use of species specific, puromycin-selected, MS5-Rh-DLL1 cells, which ensured stable and high DLL1 expression, may underlie this enhanced Notch signaling and accelerated lineage specification. Moreover, similar developmental kinetics were observed when human HSPCs were cultured in the Rh-ATO, indicating that the improved efficiency is not species restricted.

Phenotypic analyses confirmed that Rh-ATO cultures generate hallmark thymocyte intermediates, including CD34⁺CD38⁻CD7⁺ TSP-like cells, CD4⁺ ISPs, and CD4⁺CD8⁺ DPs, culminating in mature CD4⁺ and CD8⁺ T cells. The emergence of CD38⁺ single-positive subsets, analogous to recent thymic emigrants, further supported the fidelity of this platform to physiological thymopoiesis. The detection of TCRαβ-expressing cells capable of producing IFN-γ, TNF-α, and IL-2 highlights the functional maturation achieved within this system—a critical benchmark for translational application.

Earlier attempts to generate T cells from NHP HSPCs employed human or porcine thymic stromal cells (Gardner et al., 1998; Rosenzweig *et al*., 1996b; Rosenzweig *et al*., 2001), or murine OP9-DLL1 co-cultures (Radtke *et al*., 2017), While informative, these systems were limited by dependence on primary thymic tissue or incomplete recapitulation of T cell development, particularly the CD8⁺ lineage. The Rh-ATO system overcomes these barriers by providing a renewable, standardized platform that eliminates the need for primary thymic stroma while enabling full developmental progression, including functional TCR acquisition.

There are several limitations to this study. Although Rh-ATO recapitulates major aspects of thymopoiesis, we did not evaluate TCR repertoire diversity, an important metric for assessing immune breadth. Additionally, our studies focused on bone marrow–derived HSPCs; extending this approach to cord blood or mobilized peripheral blood progenitors will broaden translational utility. We also observed a reduced frequency of CD4 T cells compared with CD8 T cells using the Rh-ATO system. This is similar to the previous finding attributed to reduced MHC-II expression in MS5 system (Seet *et al*., 2017). Future work should also investigate approaches to optimize MHC-I and -II expression within the organoid microenvironment to potential a balanced selection of CD4 and CD8 T cell subsets.

In conclusion, we present the first rhesus-specific ATO platform capable of supporting complete T cell differentiation from both NHP and human HSPCs. This system faithfully models canonical thymic development and generates functionally competent T cells, providing a versatile preclinical tool to interrogate T cell biology, regenerative immunology, and immune reconstitution strategies in chronic infection and therapeutic contexts.

## Materials and Methods

### Construction of recombinant mouse bone marrow stromal cell (MS-5) expressing Rhesus macaque (Macaca mulatta) delta like canonical notch ligand 1 (DLL1)

#### Construct design

An HIV based lentiviral (LV) expression cassette was designed to express the coding sequence of *Macaca mulatta* (Rhesus macaque) delta-like canonical Notch ligand 1 (DLL1) based on reported sequence (NCBI RefSeq: XP_014993255.1). The 5’ long terminal repeat (LTR) was placed under rouse sarcoma virus (RSV) promoter replacing proviral promoter U3 region (ΔU3). Rh-DLL1 was codon optimized for expression in murine system and was placed under MNDU3 promoter. To enable fluorescent tracking, enhanced green fluorescent protein (eGFP) was co-expressed via an internal ribosome entry site (IRES). For antibiotic selection, puromycin *N*-acetyltransferase coding sequence was incorporated downstream of an mPGK promoter. Vector construction and generation of VSV-G pseudotyped particles were outsourced to VectorBuilder Inc.

#### Lentiviral transduction of MS-5 cell line

Murine bone marrow stromal MS-5 cell line (Creative Bioarray Cat #CSC-C2763) were maintained in complete DMEM supplemented with 10% fetal bovine serum (FBS; Cat #A5256701, Thermo Fisher scientific), 1% L-glutamine, 1% HEPES buffer, and 1% penicillin–streptomycin. at 37 °C in 5% CO₂. For transduction, 80% confluent MS-5 cultures were seeded at 1 × 10⁶ cells/well in six-well plates (Day 0). On Day 1, medium was replaced with 1 mL DMEM containing 5 µg/mL polybrene and LV particles encoding *nhpDLL1*. Cells were incubated overnight at 37 °C. On Day 2, viral supernatant was replaced with fresh DMEM, and cells were further incubated for 24 h. On Day 3, selection was initiated using 5 µg/mL puromycin, with medium refreshed every 48–72 h until complete selection was achieved. Recombinant MS-5-DLL1 cells were expanded and cryopreserved for downstream assays.

### Characterization of transduced MS-5 expressing Rh-DLL1

#### Validation of expression

The transduced MS5 cells was characterized for eGFP as well as surface expression of Rh-DLL1. 70%-80% confluent MS5 as well as MS5-Rh-DLL1 cell lines were trypsinized, washed with PBS, and counted. 1×10^6^ cells each from MS5 and MS5-Rh-DLL1 was transferred into 5mL FACS tubes and centrifuged at 300 x g for 5 minutes. Supernatant was discarded and cell pellet was resuspended in 100μl FACS buffer containing live/dead stain (CAT #L34975, Invitrogen) and anti-DLL1 antibody (CAT #564414, BD Biosciences). Cell and antibody mixture was incubated on ice for 20 minutes followed by two washes by FACS buffer by centrifugation at 300 x g for 5 minutes. Supernatant was discarded stained cell pellet was resuspended in 0.5mL FACS buffer. The Flowcytometry acquisition was performed to detect the expression of eGFP and Rh-DLL1. MS5 cells were used as negative control.

#### Mycoplasma control

The Rh-DLL1 cell line was tested for Mycoplasma contamination using the Universal Mycoplasma Detection Kit (ATCC, Cat. No. 30-1012K) following the manufacturer’s protocol. Cells (10⁴-10⁵) were harvested without enzymatic dissociation to preserve Mycoplasma membranes, pelleted by centrifugation (12,000 rpm, 3 min, 4°C), and lysed in 50 µL lysis buffer at 37°C for 15 minutes. A 5 µL aliquot of lysate was added to a 25µL PCR reaction containing 2× PCR Master Mix, Mycoplasma-specific primers (targeting 16S rRNA), and PCR-grade water. Reactions included positive and negative controls. PCR was run with the following conditions: 94°C for 2 min; 35 cycles of 94°C for 30 sec, 55°C for 30 sec, 72°C for 1 min; and a final extension at 72°C for 10 min. Products were resolved on a 1.5% ethidium bromide-stained agarose gel in 1× TAE buffer, run at 100–120 V for 30–40 min. A 434 bp band indicated a positive result; absence of a band indicated no contamination, with proper control validation.

### Processing of Rhesus macaque thymus tissue

Intact thymus tissue was processed to obtain thymocytes. Briefly, the thymus was sliced into small fragments using a sterile scalpel in a 10 cm² round culture dish. The tissue fragments were further minced with sterile scissors. To facilitate thymocyte release, the minced tissue was gently pressed against the culture dish using a sterile syringe plunger, followed by emulsification through repeated passage through an 18-gauge needle to disrupt larger aggregates. The resulting suspension was filtered through a 100-µm cell strainer. The filtrate was washed with PBS by centrifugation at 300 x g for 5 minutes. The supernatant was discarded, and the cell pellet was resuspended in 3 mL of ACK lysis buffer (Cat. #118-156-101, Quality Biological) and incubated for 3-4 minutes at room temperature. Following lysis, the mixture was diluted with FBS-supplemented PBS to a final volume of 25 mL. Cells were washed twice with FBS-supplemented PBS, and the final pellet was resuspended in an appropriate volume of buffer, counted, and either used immediately or stored. All steps and buffers were maintained on ice unless otherwise specified.

### Processing of Rhesus macaque peripheral blood mononuclear cells

Blood samples were collected in Vacutainer CPT™ mononuclear cell preparation tubes and centrifuged at 1,500 × g for 30 minutes. Following centrifugation, the plasma fraction was removed, and the buffy coat was transferred into a sterile 50 mL conical tube. Cells were washed with PBS by centrifugation at 300 × g for 5 minutes. The supernatant was discarded, and the pellet was resuspended in 3 mL of ACK lysis buffer and incubated for 3–4 minutes at room temperature. The cell suspension was then diluted with FBS-supplemented PBS to a final volume of 25 mL and centrifuged at 300 x g for 5 minutes. Cells were further washed with FBS-supplemented PBS as before, and the final pellet was resuspended in an appropriate volume of buffer, counted, and either used immediately or stored.

### Processing of human mobilized peripheral mononuclear cells

Human mobilized peripheral mononuclear cells were obtained from the Emory Hospital Cell and Gene Therapy Unit through established sample release procedure. These samples were left over after the patient’s treatment and were considered as discarded material. The PBMCs were centrifuged at 300 x g for 5 minutes. After discarding the supernatant, the cell pellet was treated with ACK lysis buffer (CAT # 118-156-101, Quality Biological) to remove RBCs. The cell pellet was resuspended in 3mL ACK lysis buffer and incubated for 3-4 minutes at RT. After incubation, the ACK buffer/cell mixture was topped up with FBS-supplemented PBS and adjusted the volume to 25mL and centrifuged at 300 x g for 5 minutes. Cells were washed twice with FBS-supplemented PBS as before, and final pellet was resuspended in appropriate volume of buffer, counted, and used or stored.

### Processing of Rhesus macaque bone marrow derived biospecimen

Bone marrow aspirates were obtained from young adult rhesus macaques either through the Emory Primate Center Biological Material Procurement program or through ongoing studies involving rhesus macaques with approved IACUC. The human mobilized peripheral blood was obtained through discarded product post cell therapy obtained through Emory Clinic. specimen 4-5mL bone marrow aspirate (BMA) was diluted with 2% FBS-supplemented PBS at 1 to 1 ratio and filtered with 100-micron strainer. The strained BMA was directly layered over 15mL lymphoprep (CAT #07851; STEMCELL TECHNOLOGIES) containing SepMate conical tube (CAT #85450; STEMCELL TECHNOLOGIES). Bone marrow scrap (BMS) was first resuspended in 25mL 2% FBS-supplemented PBS and filtered using 100-micron strainer. The strained BMS was further diluted to 50 mL and split into two 25mL sample each layered over 15mL lymphoprep containing regular conical tube (CAT #430921, Corning). The BMA or BMS were centrifuged for 1500 x g for 20 minutes. The buffy coat containing mononuclear cells were collected into a separate 50mL falcon tube and washed with FBS-supplemented PBS by centrifugation at 300 x g for 5 min. The cell pellet was treated with ACK lysis buffer to remove RBCs. The cell pellet was resuspended in 4mL ACK buffer and incubated for 3-4 minutes at RT. After incubation the ACK buffer/cell mixture was topped up with FBS-supplemented PBS and adjusted the volume to 30-50mL depending upon the amount of initial BMS content and centrifuged at 300 x g for 5 minutes. Cells were washed twice with FBS-supplemented PBS as before, and final pellet was resuspended in appropriate volume of buffer, counted, and used or stored. All steps and buffer used was kept chilled at all times.

### Enrichment and isolation of CD3-CD34+ HSPCs

Both rhesus macaque and human CD34+ HSPCs were first enriched using Dynabeads CD34 isolation kit (CAT #11301D, Invitrogen) as per manufacturer’s protocol. Briefly, 40×10^6^ bone marrow mononuclear cells were mixed with 100µl Dynabeads and incubated on ice for 30 minutes. Dynabead treated cells were topped up with MACS buffer (CAT #20144, STEMCELL TECHNOLOGIES) to make final volume 2mL. The tube containing bead-bound cells was placed under magnet for 2 minutes and unbound cells were discarded. This step was repeated one more time and the bead bound cells were resuspended in 100µl buffer. Next, dynabeads were detached by adding 100µl DETACHaBEAD, incubated for 45 minutes. To enhance detachment of beads, 3mL MACS buffer was added to the bead-cell mixture and vortexed. The tube was placed under magnetic field for 2 minutes and supernatant with detached cells were collected in a separate tube. To ensure all cells were collected dynabeads were further washed and subjected to magnetic field and supernatant was collected. The enriched CD34+ cells were washed with 10mL MACS buffer to remove excess DETAHaBEAD. The cells were either stored or subjected to FACS sort. For FACS sort cells were stained with live/dead, CD34, and CD3. For generation of ATO CD3-CD34+ live cells were sorted using FACS.

### Establishment of a thymus-independent artificial organoid

To culture Rh-ATO, a previously published protocol was adapted and modified (Seet *et al*., 2017). Briefly, Rh-ATO culture media was prepared by supplementing RPMI1640 (CAT #) with 3% B27 serum free media (CAT #17504001, Thermo Fisher Scientific), 1% Glutamax (CAT #35050061, Thermo Fisher Scientific), 1% penicillin-streptomycin (CAT #P4458, Thermo Fisher Scientific), 5ng/mL stem cell factor (CAT #30007100UG, PeproTech), 50μM L-ascorbic acid (CAT # A8960, Merk), 7.5ng/mL recombinant IL-7 (CAT # 200-07-2UG, PeproTech), and 7.5ng/mL FLT3 ligand (CAT #300-19-2UG, PeproTech). Appropriate number of six well plates were filled with 1mL culture media. 0.4-μm Millicell Transwell insert (CAT #PICM0RG50, Merck) were placed into the culture media containing six well plate. Next, Rh-DLL1 cell line was trypsinized, passed through 30-micron strainer to remove any cell aggregates, resuspended in culture media, and counted. 3000-7500 sorted CD3-CD34^+^ HSPCs were aggregated with rhDLL1 at 1 to 20 ratio and mixed by gently pipetting up and down few times. The cell aggregate was centrifuged at 300 x g for 3 minutes. The supernatant was carefully aspirated, and the cell pellet was resuspended in 5μl culture media per Rh-ATO. To establish Rh-ATO, 5μl of cell aggregate was carefully dropped on the insert. The plate containing cell aggregates was incubated in a humidified 37C incubator with 5% CO2. The media was replaced every 3-4 days and Rh-ATO was cultured for 1 to 12 weeks depending upon the requirement of the experiment and number of T cells needed.

### Rh-ATO phenotypic analysis by flow cytometry

To establish the T cell developmental kinetics, Rh-ATO were processed for phenotypic analysis at week 1, 2, and 4 of culture. Individual Rh-ATO was processed by pipetting 1mL FACS buffer onto the culture insert carrying the Rh-ATO and gently disrupting the aggregate using the pipette tip. Broken aggregates were pipetted in and out few times to ensure releasing lymphocytes into the solution. The lymphocyte rhDLL1 aggregate cell was filtered by passing through 100-micron strainer, and surface immunostaining was performed. To analyze T cell developmental stages cells were incubated with FACS buffer containing live/dead, CD45, CD34, CD38, CD3, CD8, CD4, CD7, TCR, CD56, CD20. The flowcytometry was performed using BD Symphony FACS analyzer and the data was analyzed using Flowjo v10.

### Rh-ATO derived CD3+ T cell enrichment and cytokine assay

CD3+ T cells were enriched from 6 Rh-ATOs at 12 weeks of culture using Pan T cell isolation kit as per manufacturer’s protocol. Briefly, Rh-ATO was processed as described above and cells were centrifuged at 300xg for 10 minutes. Supernatant was discarded and cell pellet was resuspended in 40μl of MACS buffer (Cat #20144, STEMCELL TECHNOLOGIES). 10μl of Biotin-antibody cocktail was added to cells and incubated on ice for 10 minutes. Cells were washed with 2mL MACS buffer and cell pellet was resuspended in 80μl of MACS buffer followed by addition of 20μl anti-Biotin microbeads. The mixture was incubated on ice for 15 minutes followed by washing with 2mL buffer. The cell pellet was resuspended in 500μl of buffer and subjected to magnetic separation. For magnetic separation, LS column was placed on a magnetic stand (Cat #130042401, Miltenyi Biotec). The column was prepared by passing 3mL MACS buffer followed by applying cell suspension. The flow through was collected in a sterile tube. The column was washed 3 times each with 3mL MACS buffer and flow through was collected. All steps were performed under sterile condition using chilled buffer. The collected T cell fraction was processed for stimulation with phorbol 12-myristate 13-acetate (PMA) and ionomycin (i), followed by intracellular cytokine staining as previously described (Rahman et al., 2021). Briefly, T cells were seeded in 200μl complete RPMI containing either PMA (80 ng/ml) plus i (1 μg/ml) or without PMAi. The mixture was incubated at 37C, 5% CO2 for 2h followed by addition of Glogi-stop (CAT #51-2092KZ, BD Biosciences) and Golgi-plug (CAT #512301KZ, BD Biosciences) and further incubation for 4h. Intracellular cell cytokine staining: PMAi stimulated and unstimulated cells were washed with FACS buffer and incubated with 100μl FACS buffer containing anti-CD3, IFNγ, TNFα, and IL-2 antibodies. The cell and antibody cocktail were incubated for 20 minutes on ice followed by washing with 500 FACS buffer twice. Upon 2^nd^ wash the supernatant was discarded and cell pellet was resuspended in 200μl BD Cytofix/Cytoperm fixation and permeabilization solution (CAT # 554722, BD Biosciences) per test and incubated for 20 minutes on ice. The cells were washed twice with Perm wash buffer and further incubated with intracellular stain in 100μl perm buffer containing anti-IFNγ, TNFα, and IL-2 antibodies. The cell antibody mix was incubated for 20 minutes on ice followed by two washed with 500μl perm wash buffer (Cat #BDB554723; BD Biosciences) and a final wash with FACS buffer. The cells were resuspended in 200μl FACS buffer and flowcytometry was performed.

### Antibodies

Antibodies used in this study are as follows: CD3 (clone SP-34-2; BD Biosciences), CD45 (clone D058-1283 <u>(RUO)</u>, BD Biosciences), CD8 ( clone SK1; BD Bioscience), CD4 (clone L200; BD Biosciences), CD34 (clone 561, BioLegend), CD38 (clone AT1), CD7 (clone M-T70, BD Biosciences), TNFα (MAb11; BD Biosciences), IL-2 (MQ1-17H12; BD Biosciences). IFNγ (B27; BD Biosciences), CD20 (clone 2H7, BioLegend), CD56 (clone NCAM162, BD Biosciences), TCRαβ (clone R73, BioLegend).

### Quantification and statistical analysis

Statistical significance was calculated using a two-tailed unpaired nonparametric Mann-Whitney test. GraphPad Prism version 8.4.3 (GraphPad Software) was used to perform data analysis and statistical tests.

## ACKNOWLEDGMENTS

We thank Stephanie Ehnert, Stacey Weissman, and other member of the research tech services team at the Emory National Primate Research Center (ENPRC). We thank Jennifer S. Wood, Rachelle Lauren Stammen, Sherrie Jean and all current and former veterinarian scientists at the ENPRC. We thank Elizabeth H. Curran and members of the necropsy team at the ENPRC. We thank Kiran Gill, Ankur Saini, Robert E. Karaffa II, Kametha Fife, Sommer Durham, and members of the flowcytometry cores at the ENPRC and Emory Vaccine Center, Emory University. We thank the Kay Lee Summerville and members of the ENPRC Biological Material Procurement program. BioRender was used to make the schematic shown in Figure 2A. We thank Prof. Gay M. Crook, Eli and Edythe Broad Center of Regenerative Medicine and Stem Cell Research, DGSOM, UCLA, for helpful advice in setting up the system. The content is solely the responsibility of the authors and does not necessarily reflect the official views of the National Institute of Health.

## AUTHOR CONTRIBUTIONS

S.A.R. conceived and designed the research, brought funding, conducted the research, analyzed data, interpreted results, and wrote the manuscript. C.W., and S.A. designed and conducted the research, analyzed the data. J.D.R. provided human PBMCs. S.B., Y.G.P., J.D.R., R.R.A. and R.P.J. helped with data interpretations. S.A.R. supervised the research. All authors read and agreed to the published version of the manuscript.

## DATA AVAILABILITY

The raw data were generated at the Emory University and will be available upon request from the corresponding author.

